# Time cells in the human hippocampus and entorhinal cortex support episodic memory

**DOI:** 10.1101/2020.02.03.932749

**Authors:** Gray Umbach, Pranish Kantak, Joshua Jacobs, Michael Kahana, Brad E. Pfeiffer, Michael Sperling, Bradley Lega

## Abstract

The organization of temporal information is critical for the encoding and retrieval of episodic memories. In the rodent hippocampus and entorhinal cortex, recent evidence suggests that temporal information is encoded by a population of “time cells.” We identify time cells in humans using intracranial microelectrode recordings obtained from 27 human epilepsy patients who performed an episodic memory task. We show that time cell activity predicts the temporal organization of episodic memories. A significant portion of these cells exhibits phase precession, a key phenomenon not previously seen in human recordings. These findings establish a cellular mechanism for the representation of temporal information in the human brain needed to form episodic memories.

## Main Text

Episodic memory describes our ability to weave temporally contiguous elements into recollections of rich and coherent experiences. The activity of time cells in the mesial temporal lobe may provide a mechanism for the coding of temporal information that is necessary for the formation of these memories *(1-6)*. The spiking of these cells reliably increases at specific moments within a fixed interval, with different groups of cells tuned to represent distinct but overlapping moments. The detailed temporal information available from a population of these time cells allows the hippocampus to impose temporal order on representations of individual items.

With properties analogous to place cells, time cells offer a possible unifying mechanism for the representation of both spatial and temporal context in the hippocampus. In rodents, the activity of both time cells *(1, 4)* and place cells *(7)* predicts memory performance, and they both exhibit phase precession *(1)*, a property that provides a direct mechanism for linking together spatial and temporal information into a continuously evolving contextual representation. Subsequent experiments in rodents identified a distinct population of temporally sensitive cells in the lateral entorhinal cortex (LEC) whose activity gradually rises or decays across a given time interval (termed “ramping cells”) *(8)*. As these cells are sensitive to contextual changes during experience, they may represent the slowly evolving nature of contextual information. As such, ramping cells complement hippocampal time cells or may even modulate their activity *(8-10)*, analogous to hexadirectional grid cells in spatial navigation *(11-13)*. The characterization of time cells in rodents stimulated widespread interest among memory theorists, reflected in proposed models of episodic memory in which time cells play a critical role *(9, 14)*. To date however, time cells have not been reported in humans, and it is unknown if they support episodic memory.

To investigate whether time cells exist in humans and their potential relationship to episodic memory, we used microelectrode recordings (Fig. S1) obtained in 27 human participants who performed the free recall task (Fig. 1A), a standard assay of episodic memory. The structure of the task, with consistent and well-defined time segments for each item encoding list, facilitated our ability to detect time cells. In the task, participants studied lists of common nouns presented regularly over the course of a 1.6 second encoding interval. Following an arithmetic distractor task, participants freely recalled as many words as they could remember. They completed an average of 15.3 item lists, successfully remembering an average of 24.2 ± 11.6% of the memory items. Behavioral performance during these sessions exhibited expected patterns, including greater recall accuracy for the initial list items (a primacy effect, Fig. 1E and 1F).

**Fig. 1.**
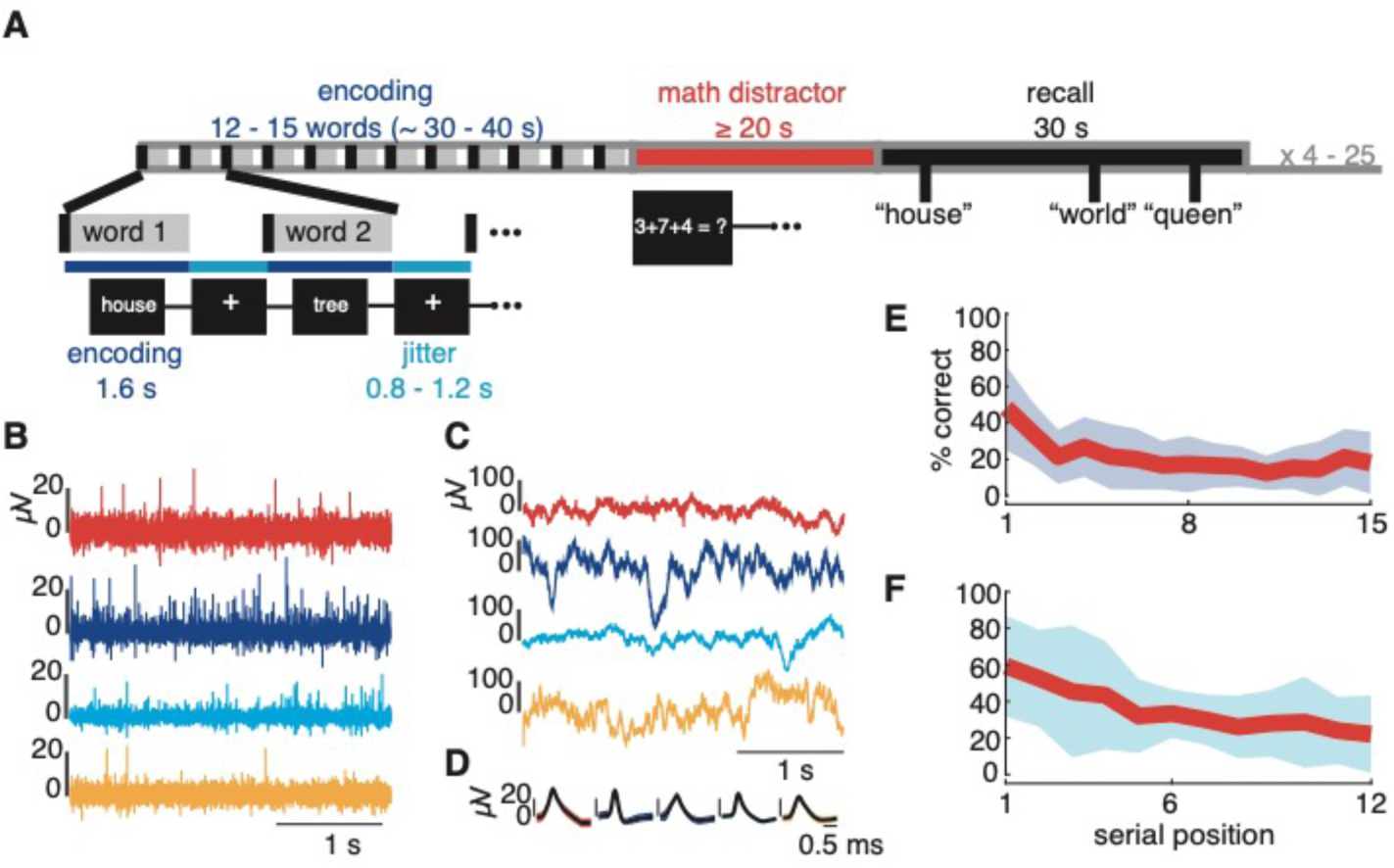
Task and performance. (A) Free recall task. (B) Noise-subtracted and band-passed (300-1,000 Hz) signal from 4 different channels used for single-unit isolation. (C) Noise-subtracted and low-passed (<300 Hz) signal. (D) Mean waveforms of time cells extracted from the channels displayed in (B). (E) Recall performance by encoding list serial position for all recording sessions with 15-word lists (n = 18). (F) The same as (D) but for all recording sessions with 12-word lists (n = 6).

We isolated a total of 768 single units from the implanted microelectrodes (Fig. 1B-D), calculating standard quality metrics that were comparable to previously published data describing human medial temporal lobe (MTL) neurons (Table S1, Fig. S2) *(15)*. We focused our analysis on putative pyramidal neurons isolated from the hippocampus and entorhinal cortex (n = 458 and 51 respectively from each region, defined by spike width and firing rate, see supplementary materials). To represent temporal information, time cells fire within a preferred time window relative to the beginning and end of a period of fixed duration, which in our experiment was the encoding list in a free recall task. Stated another way, the firing rate of time cells is significantly modulated by the factor of time (Fig. S3). We used a non-parametric method (P < 0.05, Kruskal-Wallis test) with an associated circular shuffle procedure to identify this kind of firing pattern (79 out of 509 candidate single units met these criteria, P < 0.001, binomial test). Example time cells, exhibiting characteristic firing with a preferred temporal window are shown in Fig. 2 and S4A. We tested for the significance of time cells at the subject level as well using a shuffle procedure to test the number of time cells observed within each subject against a null distribution. This was also significant (t(25) = 4.44, p < 0.001). We observed time cells in 24 of 26 subjects in which we isolated at least one pyramidal cell. These time cells had specific and concise firing fields across the encoding list that remained consistent across lists (Fig S5B), and time cell identification was robust to alterations in parameter selection (such as time bin width, see supplementary material).

**Fig. 2.**
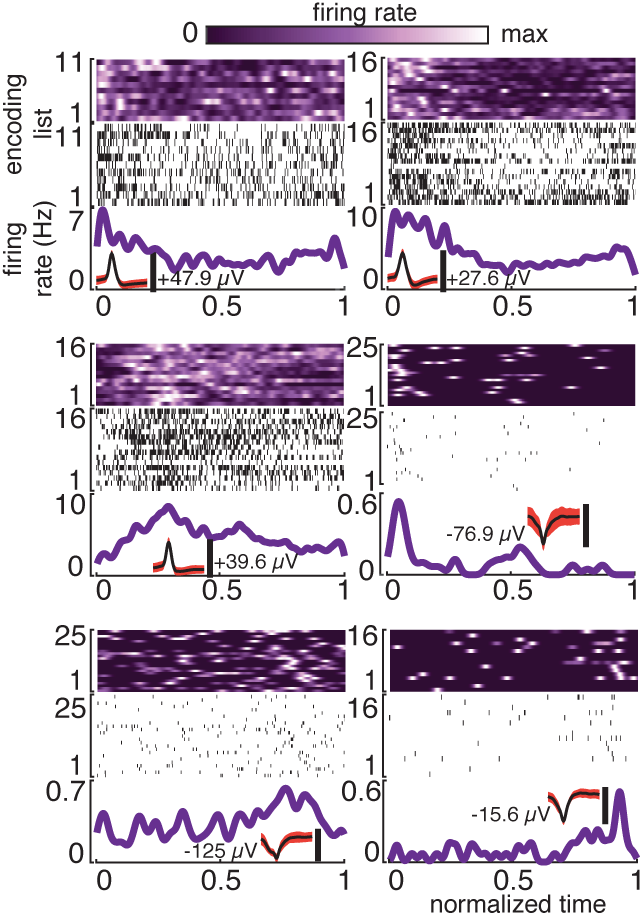
Human time cells are active during a memory task. 6 examples of time cells. Spike heat map (top), spike raster (middle), and PSTH (bottom) plotted against normalized encoding list time. The mean spike waveforms are inset.

In an approach complementary to our initial method, we implemented a recently published technique to identify time cells through modeling the firing rate curves *(6, 16)*. This method also yielded a significant number of time cells (69 of 509 candidate units, P < 0.001, binomial test, examples shown in Fig. S4C and S6). We also conducted a series of control analyses to ensure that the identification of time cells was not affected by factors such as onset of memory items within a list, the semantic content of the items, or recall success (Fig. S5, S7, and S8).

A key property of time cells is that, in aggregate, they should represent time throughout a given epoch, analogous to place cells. Across all participants, this is indeed what we observed by plotting the peri-stimulus time histograms (PSTHs) according to each cell’s preferred temporal window (Fig. 3A). Similar to previous reports of time cells in rodents, we observed denser temporal representation at early and late epochs *(6, 16)*. Time cells in the entorhinal cortex preferentially represented earlier time bins as compared to time cells recorded from the hippocampus (χ^2^(1) = 12.3, P < 0.001, Fig 3B).

**Fig. 3.**
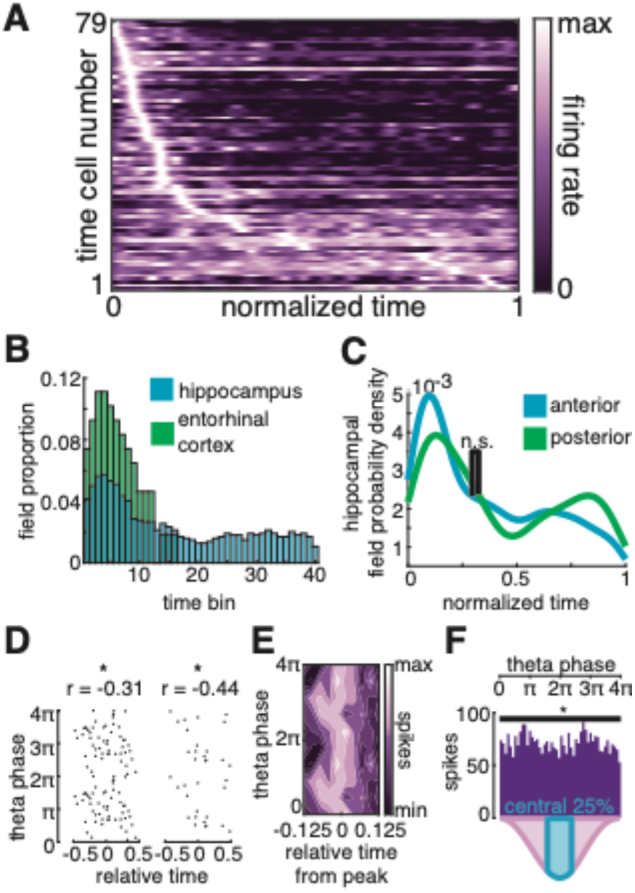
Properties of the time cell population. (A) Time cell firing rate heat map with rows organized by the time of the peak in the PSTH. (B) Histogram of the time bins covered by the time cells’ time fields. (C) Comparison of the density of time fields represented by time cells in the anterior and posterior hippocampus. (D) Phase-time plots for two example time cells, in their preferred time field. The circular-linear correlation coefficient value is shown above. (E) Heat map of spike counts obtained by superimposing the central 25% (centered on the time of peak time cell activity) of the phase-time plots from all time cells that demonstrated significant precession. (F) Phase histogram of spikes falling within the central 25% of the time fields of all time cells demonstrating precession. For all plots, n.s. = not significant, *P < 0.05.

We asked whether humans additionally exhibit ramping cells, defined as cells whose activity increases or decreases across the duration of a temporal epoch *(8)*. These may provide complementary information to time cells. Following methods established in recent rodent publications, we identified 82 ramping cells (Fig. S4B and S9, P < 0.001, binomial test). Consistent with the initial description of ramping cells in rodents, we observed that ramping cells occur more frequently in the EC as compared to hippocampus (17/51 EC units versus 65/458 hippocampal units, χ^2^(1) = 12.4, P < 0.001).

Given the proposed similarities between time cells and place cells *(17)*, we asked whether time cells exhibit phase precession. This question is important for two reasons. First, phase precession provides a possible mechanism for integrating and organizing events in time. Second, precession has not previously been demonstrated in human microelectrode recordings. We implemented established methods for measuring phase precession *(18)*, focusing on the firing of time cells within their preferred time fields. We measured theta phase angle for all spike events centered at when time cells exhibited their highest firing rate (the middle of the time field). We evaluated precession within the 1-10 Hz range, encompassing frequencies that exhibit mnemonically relevant properties in humans such as phase locking, phase reset, and power increases during successful memory encoding *(19, 20)*. Twenty-four time cells demonstrated a significant correlation between time and phase at one or more of these frequencies (Fig. 3D and 3E, Fig. S10, P < 0.001, binomial test). Similar to theta precession described in rodents for place cells *(21)*, peak firing within the time field occurred near the trough of the theta cycle (Fig. 3F, 95% confidence interval for phase in the central 25% of the time field: 110 deg – 197 deg, P = 0.018, Rayleigh test). These time cells had a mean precession rate of 69 ± 48 degrees/s and a median correlation of -0.30 between phase and time.

A unique feature of our data set compared to previous human microelectrode experiments was the availability of recordings from both the anterior and posterior hippocampus. We isolated 34 time cells from the anterior hippocampus and 22 from the posterior hippocampus. These recordings provided an opportunity to compare the width of time fields for anterior versus posterior hippocampal time cells, motivated by the well-known finding that rodent place cells exhibit wider fields in the ventral (analogous to anterior) compared to dorsal hippocampus *(22)*. However, we observed no significant difference in time field duration according to hippocampal longitudinal location (Fig. S11, median duration 2.72 s versus 2.59 s respectively, z(177) = 1.21, P > 0.2), nor did time field location differ between anterior and posterior time cells (z(50) = - 1.05, P > 0.2, Fig 3C).

We analyzed the relationship between time cell activity and memory behavior, asking whether firing patterns of time cells are associated with the tendency of human subjects to remember items presented adjacent to one another in time during encoding (quantified using the temporal clustering factor or TCF) *(23)*. This is because the temporal clustering of memory items relies directly on temporal contextual information, and time cells potentially provide this information. In rodents, variation of time cell firing relative to a preferred time field predicts future memory errors *(1)*, and we therefore hypothesized that this same type of variability would predict decreased temporal clustering behavior. We reasoned that less precision in the signal provided by time cells would decrease participants’ ability to temporally cluster items. We used an estimate of time cell firing consistency calculated as the average similarity between a time cell’s firing pattern on a single item list and its average firing pattern across all lists. We termed this parameter ρ (Fig. 4A and 4B, see supplementary material). Higher ρ means that a cell’s firing pattern remains consistent across encoding lists, while lower ρ indicates greater variability in firing patterns across lists. Partitioning time cells into those with high and low ρ values (split at the median), we found that those with higher ρ are associated with greater temporal clustering of items by subjects at the time of memory retrieval (z(77) = 2.56, P = 0.011, rank sum test, Fig. 4A-C). This finding supports a link between the fidelity of temporal information provided by time cells and the tendency of participants to use temporal information when retrieving memory items. We did not find an association between ρ and semantic clustering (z(77) = -1.21, P > 0.2), indicating that consistency of time cell firing specifically predicts the temporal organization of memories. Further, consistent temporal representation provided by time cells (higher ρ values) increased the likelihood that individuals would recall memory items located within a time cell’s preferred field (Fig. 4E, z(70) = 2.23, P = 0.025, rank sum test), although ρ did not significantly predict overall performance (Fig. 4D, z(77) = -1.93, P = 0.054, rank sum test). This latter result likely reflects the fact that several variables beyond temporal contextual information affect performance in the free recall task. We also observed a more general connection between time cell activity and memory behavior, namely that time cells appear to provide more detailed information near the beginning and end of an item list, especially during the time segments in which the first few memory items are presented (supplementary text, Fig. S12).

**Fig. 4.**
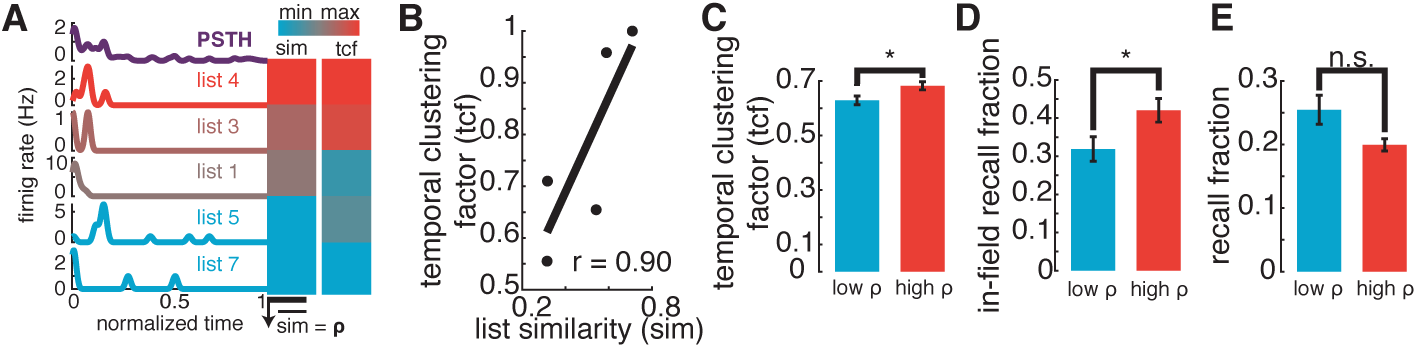
Time cell stability promotes temporal clustering and influences which serial positions are recalled. (A) Single cell example of the correlation between time cell firing pattern similarity (sim) and temporal clustering (TCF). (B) Scatter-plot demonstrating a positive association between sim and tcf for the example neuron from (A). (C) Comparison of session-level temporal clustering between cells with high and low ρ. (D) Comparison of in-field memory performance between cells with high and low ρ. (E) Comparison of memory performance between cells with high and low ρ. For all plots, n.s. = not significant, *P < 0.05, bars represent mean values and error bars the standard error of the mean.

Our findings connecting the consistency of time cell firing and temporal clustering behavior explicate a core feature of human memory and potentially unlock new strategies for neuromodulation. In contrast to the direct representation of temporal context provided by these cells, previous data in humans have utilized aggregate measures of temporal lobe single units and local field potential recordings to show that neurophysiological states drift slowly over longer time scales *(23-25)*. This drift in activity may provide coarse temporal information. The relationship between these aggregate estimates of activity and the precise information provided by time cells remains a key area for further investigation. In this regard, ramping cells specifically may connect representations at different time scales by modulating hippocampal time cell activity *(9, 10, 26)*. Taken together, our findings and these previous human studies support the contention that the representation of time in the brain arises inherently from continuously evolving brain states *(8, 14)*, although alternate memory paradigms are needed to show that human ramping cells flexibly respond to different temporal epochs *(8)*. Data in rodents and primates suggest that variability in place cell firing (analogous to our ρ measurement) reflects the representation of item and contextual features beyond spatial information *(27, 28)*, suggesting that the spiking activity of subpopulations of time cells may encode item-level information such as semantic category or reward conditions, depending upon the task. A complete understanding of time cells in humans will ultimately require identifying what additional information may be encoded by these cells and how they respond to different task demands *(16, 29)*.

The temporal organization of memories constitutes a key function of the medial temporal lobe *(30)*. The discovery of time cells that we report establishes a direct neurophysiological mechanism for the representation of temporal contextual information that is necessary for this function. The demonstration of time cells in humans also lends critical support to prominent models of memory that have posited temporal coding mechanisms as central to the process of associative memory *(31, 32)*. Finally, the possible modulation or restoration of information provided by time cells represents an intriguing strategy to treat episodic memory deficits due to conditions such as traumatic brain injury and Alzheimer’s disease.

## Acknowledgments

We thank G. Konopka and Z. Tiganj for helpful remarks on the manuscript. We are grateful for all the patients who volunteered to participate in this study.

## Funding

This work was funded by National Institutes of Health (NIH) research project grants to B. L. (R01-NS107357), J. J. (R01-MH104606), and M. K. (R01-MH55687), as well as by support from the Southwestern Medical Foundation to B. E. P.

## Authors contributions

G. U. and B. L. conceptualized the project. G. U., P. K, and M. S. curated the data. G. U. performed the analyses and visualization and developed the required software. B. L. acquired the necessary resources and financial support as well as managed and supervised the project. P. K., M. S., and B. L. conducted the investigation. G. U., J. J., B. E. P., M. K., and B. L. developed the methodology. G. U. and B. L. validated the results and wrote the original manuscript. G. U., J. J., B. E. P., M. S., M. K., and B. L. reviewed and edited the final manuscript.

## Competing interests

The authors declare no competing interests.

## Data and materials availability

All data and software used to analyze the data are available upon request from the corresponding author. Data and software will be additionally available to the public at https://www.utsouthwestern.edu/labs/tcm/.

## Supplementary Materials

## Materials and Methods

### Subjects and Electrophysiology

We consented a total of 27 human epilepsy patients undergoing clinical seizure mapping at either Thomas Jefferson University Hospital (TJUH) or University of Texas Southwestern Medical Center (UTSW) for enrollment in the study. The 27 subjects completed a total of 40 sessions of the memory task. We treated each session as independent for purposes of statistical analyses unless otherwise specified. Patients were implanted with depth electrodes at locations dictated by clinical interest. Electrodes featured both macroelectrode contacts along their length and 9 x 40 um platinum-iridium microwires extruding from the tip. We sampled the broadband signal at either 30 kHz with NeuroPort (Blackrock Microsystems, UTSW) or 32.6 kHz with Cheetah (Neuralynx, TJUH) recording systems. We localized the microwires by co-registering post-implantation computed tomography (CT) scans with pre-operative magnetic resonance imaging (MRI). Neuroradiologists then confirmed localizations. We used BrainNet Viewer *(33)* to plot electrode localizations.

Before spike detection and sorting, we bandpass filtered the broadband signal from 300 to 1,000 Hz and cleaned it with a volume conduction subtraction algorithm *(34)*. We then used Combinato *(35)* for spike detection and clustering. Subsequently, putative units were manually reviewed by the first author for the (1) shape of the mean spike waveform, (2) number of inter-spike intervals (ISI) below a threshold of 3 ms, (3) shape of the ISI distribution (4) stationarity of unit spiking, and (5) similarity to other mean spike waveforms isolated from the same microwire. Based on these criteria, we either merged or discarded putative units. For all microwires with more than one isolated unit, we calculated pairwise isolation distances *(36)* to examine the degree of separation of one cluster from other clusters. We isolated a total of 768 units (also referred to as “cells” and “neurons” throughout the manuscript). Only putative pyramidal cells located in either the entorhinal cortex or hippocampus (509/768) were considered for all time cell analyses. We identified these as units with a firing rate of less than 5 Hz *(37-39)* and with a “wide” (> 0.40 ms) spike width *(40)*. We defined the spike width as the trough-to-peak time *(18)*. Hartigan’s DIP test confirmed bimodality of the spike width distribution (P < 0.001) *(18)*. We then fit a fourth order polynomial to the data and used the spike width at the local minimum that split the two modes of the distribution (0.40 ms) to divide units into those with “narrow” and “wide” spike widths.

For all 768 units isolated, we computed quality metrics as reported in a similar, large human single unit dataset *(18)*. Units isolated per channel, mean firing rate, percentage of inter-spike intervals less than 3 ms, modified coefficient of variation (CV2), signal-to-noise ratio (SNR) of both the peak of the mean waveform and the mean of the mean waveform, and isolation distance distributions were all comparable to this reference high quality dataset. These data are displayed in Table S1 and Fig. S2.

When the phase of the theta local field potential (LFP) was required for an analysis, we first lowpass filtered the signal at 300 Hz. We then applied a notch filter at 60 Hz. Finally, we down-sampled the signal by a factor of 30 or 32, depending on the original sampling rate. Phase was then extracted by convolving 6 evenly log-spaced Morlet wavelets, all with a width of 6, at frequencies between 1.68 and 9.51 Hz.

### Behavioral Task

Subjects completed a free recall task, delivered at clinical bedside via a laptop computer. The task consisted of a repeating encoding-distractor-retrieval paradigm. During the encoding section, patients studied a sequence of words that appeared on the screen. Each word stayed on screen for 1.6 s. A 0.8-1.2 s jitter period separated neighboring words. TJUH subjects studied 15-word lists. UTSW subjects studied 12-word lists. During the distractor period, participants completed simple arithmetic problems for at least 20 s. All problems followed the format of A + B + C = ?. Following this, subjects entered into a 30 s retrieval phase, during which they recalled as many words as able from the most recently studied word list. Subjects were allowed breaks in between word lists. Subjects repeated this sequence for as many lists as they could, up to 25. For all analyses save the performance versus serial position behavioral analysis, sessions with 15-word lists and 12-word lists were considered together. We rejected sessions with seizure activity or aura.

### Time Cell Identification

We fit three models to the spike train data to identify time cells, field cells, and ramping cells. Each method was rooted in previous place or time cell studies *(6, 8, 19, 41)*. For the first method *(41)*, we convolved the session-wide spike train of each cell with a Gaussian kernel with a standard deviation of 0.5 s in order to obtain an instantaneous firing rate at every sample in the recording. We refer to the convoluted spike train as the “tuning curve.” We then isolated the sections of the tuning curve corresponding to each of the encoding lists. These list-level tuning curves were then aligned by normalizing the time domain to account for small differences in absolute time from list to list due to inter-word jitter. Next, we divided the encoding list tuning curves into 40 equally spaced, normalized-time bins (ranging in absolute time from roughly 0.75-1 s, depending on jitter and number of words on the list), and took the mean of the firing rate at all samples within each bin. After constructing a list-by-time-bin matrix, we tested for temporal modulation of firing rate by normalized-time bin with a Kruskal-Wallis test. We chose a mean rank-based test to limit the influence of a single instance of a high firing rate in a given time bin and because firing rate is Poisson, not normally, distributed. We labeled cells with a P_actual_ < 0.05 with this procedure as potential time cells and passed them on to the next step of testing. We then subjected all potential time cells to permutation testing. To do this, we randomly circularly shuffled the session-wide tuning curve and then repeated the described procedure. We did this 1,000 times for each cell. P_shuffle_ was finally obtained by comparing P_actual_ to the distribution of P-values obtained from the permutation procedure. We defined P_shuffle_ as the number of shuffles producing a smaller P-value than P_actual_ divided by 1,000. Only potential time cells with P_shuffle_ < 0.05 were ultimately considered “time cells.” We tested the fraction of cells determined to be time cells (79/509) for significance with a one-sided binomial test, with a false discovery rate (FDR) of 0.05. Note that this model did not assume a specific relationship between time and firing rate, only that time modulates firing rate. Practically, this means that time cells are not required to have a single temporal field of increased activity. For this cohort of time cells, we defined “time fields” as any consecutive time window during which the cross-encoding list mean tuning curve (the peri-stimulus time histogram, PSTH) was above the session-wide mean firing rate of that cell for at least the duration of one of the time bins used in the Kruskal-Wallis. To increase the temporal specificity of time fields, we used 400 time bins (ranging in absolute time from roughly 0.075-0.1 s, depending on jitter and the number of words on the list) to generate the PSTH. A minority (7/79) of time cells did not have a time field by this definition. We defined the time field center as the time point at which the time cell’s firing rate reached its maximum within the field. For each anterior and posterior hippocampal time cell, we calculated the center of mass of all its time fields in order compare the temporal representation of time cells along the hippocampal longitudinal axis.

### Time Field Cell Identification

We next identified time cells with a single time field, referred to as “field time cells” or “field cells.” To do so, we utilized maximum likelihood estimation methods, adapting a previously published procedure *(6, 19)*. We first down sampled the spike train by a factor of 32 or 30, depending on the original sampling rate. We then compared the fits of two models that describe the likelihood of spiking activity at any given sample along the length of the encoding list. The time field model was specified by a total of four parameters, included a Gaussian field of increased firing probability located somewhere along the length of the encoding list:

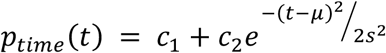

where “p” is the probability of a spike at time “t.” “t” in the equation is normalized time. c_1_ and c_2_ are constants assigning the influence of the constant and Gaussian term to the overall probability of spiking. So that probability did not exceed 1, bounds were set so that c_1_ + c_2_ ≤ 1. “μ” and “s” are the mean and standard deviation of the Gaussian field respectively. The former was bound between 0 and 1, so that the mean of the field was located within the encoding list. To prevent excessively large Gaussian fields appearing as a flat line across the list, the standard deviation was bound at 1/6. This way, at largest, the field length would approximate the encoding list length. A constant model assumed that the probability of spiking is stationary across the encoding list:

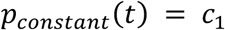

We used matlab’s *particleswarm* with *fmincon* as a hybrid function to minimize the negative log-likelihood of these models to solve for their parameters. Negative log-likelihood was specified by the equation below:

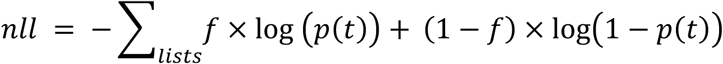

where “f” is the actual spike train at all time points in the encoding list and p(t) is as defined by either p_time_ or p_constant_. Superiority of the time model was determined by comparing log-likelihood values from the two models with matlab’s *lratiotest*. We fit the model to data from all lists, only odd lists, and only even lists to avoid a single list driving the effect. To be labeled a field cell, for all encoding list partitions, the time field model must have a significantly higher log-likelihood than the constant model (P < 0.05). Additionally, to force consistency in odd and even lists, the time fields predicted by odd and even list models must overlap (|μ_even_ – μ_odd_| ≤ s_even_+ s_odd_). We tested the fraction of field cells identified (69/509) for significance with a one-sided binomial test with an FDR of 0.05.

### Ramping Cell Identification

We finally searched for ramping cells with a methodology similar to that previously described *(8)*. To reduce the dimensionality of the data, we calculated firing rate by counting the number of spikes in 1 s time bins. All time bins corresponding to each of the encoding lists were then concatenated in series and used as the outcome variable in a generalized linear model (GLM). We built two nested GLMs, one with only a constant term, and the other with the addition of encoding list time as a predictor. We used matlab’s *fitglm* function with the distribution set to “poisson” to build the model. We utilized a log-link function, assuming an exponential relationship between time and firing rate. We considered encoding lists in three partitions: odd lists, even lists, and all lists to prevent activity on a single list from driving a statistically significant relationship. In order to be considered a ramping cell, the time predictor model had to outperform the constant model for all three list partitions. We said the time model outperformed the constant model if (1) the log-likelihood for the time model was significantly greater than that of the constant model (P < 0.05, using matlab’s *lratiotest*), (2) the time model better explained the data than the constant model by the Chi-square goodness-of-fit test (P < 0.05), and (3) the coefficient describing the relationship between firing rate and time was significantly different from zero by a two-tailed t-test (P < 0.05). We tested the fraction of cells meeting these criteria (82/509) for significance with a one-sided binomial test, with an FDR of 0.05. No R^2^ threshold was used in the initial identification procedure.

### Time Cell Control Analyses

To increase confidence in our ability to identify an above-chance number of time cells, we employed a secondary statistical significance procedure. For this, we circularly shuffled the spike trains of all of the subject’s neurons and performed the Kruskal-Wallis based time cell identification procedure. We repeated this 1,000 times for each subject, counting the number of time cells isolated for each shuffle. Using this distribution of time cell counts, we transformed the actual number of time cells obtained from each subject into a z-score. We then ran a one-sample, two-tailed t-test against zero on of the distribution subject z-scores to see if the number of time cells obtained significantly differed from chance.

We conducted a series of analyses to disambiguate time’s influence on firing rate from the covariates of stimulus onset, recall success, and semantic information of the studied words. To do this, we built a GLM with firing rate as the outcome and with four categorical variables plus a constant term as the predictors. We fit the GLM with matlab’s *fitglm* tool, with the distribution set to “poisson,” using a log-link function. We calculated firing rate by counting spikes in 1 s bins across each of the encoding lists. We set the first categorical predictor, stimulus onset, to 1 when the time bin corresponded to the bin of word onset on the screen, and 0 otherwise. The second, recall success, we set equal to 1 for bins during which a word that was later recalled was displayed, and 0 otherwise. The third, semantic information, was equal to 1 when in a time bin covered by the ideal “semantic field” (see Latent Semantic Analysis), and 0 otherwise. Similarly, the fourth, time, we defined as 1 when in a time bin covered by the peak time field, and 0 otherwise. When a cell had no time field, these latter two predictors were monotonically 0. We fit the GLM to two cohorts of cells: all time cells and all subsequent memory effect (SME) cells that were not concurrently time cells (see Subsequent memory effect cell identification). A significant relationship between a given covariate and firing rate was considered a positive coefficient significantly different than zero by a two-tailed t-test. The numbers of cells from each group significantly related to each of the covariates were counted and compared.

Additionally, for all time cells, we compared the firing rate during all successful encoding events within that cell’s time field to the firing rate during all unsuccessful encoding events within the time field. This was done at the cell level, by averaging across in-field encoding events, subtracting the average for unsuccessful encoding events from the average for successful encoding events, and comparing the distribution to zero with a one-sample, two-tailed rank sum test (Fig. S5B).

### Subsequent Memory Effect Cell Identification

We looked for cells demonstrating an SME by counting the number of spikes during all successful (word later recalled) and unsuccessful (word not later recalled) encoding events throughout the recording session. We then compared counts with a two-sample, one-tailed rank sum test. Cells with a P < 0.05 were considered “SME cells.” All isolated cells were considered in this analysis. We tested the fraction of identified cells (101/768) for significance with a one-sided binomial test with an FDR of 0.05. We assessed the association between SME and time cells using Yule’s Q. To obtain an appropriate test statistic, we used the t-approximation:

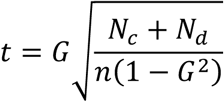

where G is Yule’s Q, N_c_ is the number of concordant pairs, defined as the number of cells that are both time and SME cells multiplied by the number of cells that are neither time nor SME cells, N_d_ is the number of discordant pairs, defined as the number of time cells that are not SME cells multiplied by the number of cells that are SME cells but are not time cell, and n is the number of candidate neurons.

### Latent Semantic Analysis

We used a subset of 100,000 documents from a publicly available corpus of all English-language Wikipedia articles as of April 2010 *(42)* to build a word count matrix for latent semantic analysis (LSA). All occurrences of the 308 words used in the experimental task were counted across the 100,000 articles. We then used matlab’s *svd* function with the ‘econ’ option for singular value decomposition. The result of this transformation is as follows:

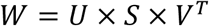

where W is the word count matrix, U contains the left singular vectors, S is a diagonal matrix containing the singular values for each semantic dimension, and V contains the right singular vectors. We multiplied U with S to generate a word-by-dimension matrix whose values communicate the semantic location of the index word in the index semantic dimension. To compare semantic information of two words, we calculated the cosine similarity between two rows of this matrix *(43)*. To control the temporal clustering results, we computed a semantic clustering factor using all word transitions across a given session. This calculation mirrored that used for temporal clustering. For each recall event, we computed the semantic similarity between the words comprising all possible word transitions. We then assigned a semantic transition score based on where the actual transition ranked among all possible transitions. We averaged all transition scores across the experiment to obtain a single value for each session.

We additionally used this analysis to add a categorical predictor communicating semantic information to the control GLM. To do so, we first calculated the pairwise similarity between all words covered by each cell’s preferred time field across all lists. We next compared this distribution of in-field similarity values to a distribution of all similarity values generated from all pairwise combinations of all words seen throughout the entire recording session with a two-sample, one-tailed rank sum test. A P_actual_ < 0.05 in this context means that the cohort of words seen while in the time field were significantly more semantically similar than were all words seen throughout the experiment. However, to control for increased similarity in a subset of words by chance random allocation, we then searched for the optimal location for a hypothetical “semantic field” of the same size as the time field. We did this by shifting the time field over all possible temporal lags until the field either began at the same time as the encoding list or ended at the same time as the encoding list. The aforementioned statistical procedure was repeated at all possible lags. We used the time field at the shift that generated the lowest P-value as the “semantic field” in the GLM control analysis.

### Precession

We calculated precession on the all-list, all-spikes level within the peak time field of each time cell. We used circular-linear correlation methods, as previously described *(21, 44).* We did this by fitting the following non-linear model, relating theta phase to time:

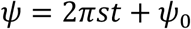

where “ψ” is the circular variable theta-phase, “s” is the slope describing the relationship, “t” is the linear variable time, and “ψ_0_” is the phase offset. We fit this model by minimizing the length of the mean resultant, given by the following equation, with matlab’s *fminbnd*:

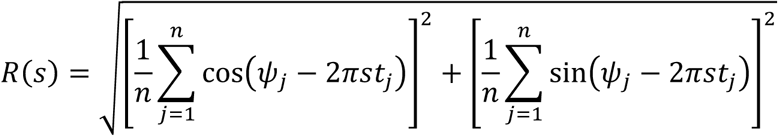

where “n” represents the number of in-field spikes. The solver was bound such that the slope could only range from 0 to ± 4π *(45)*. A time field was said to be precessing if the slope relating phase to time was less than zero. However, a field was said to demonstrate significant precession on the basis of the circular-circular correlation coefficient obtained after transforming time into a circular variable *(21, 46)*. Due to the heterogeneity of human theta, we repeated the above procedure at each of six-log spaced frequencies within the theta band for each time cell.

We then FDR-corrected the resulting vector of p-values obtained for each cell across frequencies with a Q of 0.20. We then tested the fraction of surviving cells against chance with a one-sided binomial test with an FDR of Q/2 to account for the directionality of the slope. We tested phase distributions against the null hypothesis of uniformity with a Rayleigh test. We computed all circular statistics with the matlab toolbox developed by Berens et al. *(46)*.

### Behavioral Analyses

We used the mean list-to-mean tuning curve correlation, ρ, as the predictor variable in the behavioral analyses. We calculated this by correlating all the individual list tuning curves with the cross-list mean tuning curve (PSTH) and averaging the resulting Spearman rank correlation coefficients. Thus, this metric is a measure of the consistency in the temporal modulation of firing rate from list to list. We refer to this idea as “consistency” or “stability” in the time field representation. When two-group testing was used with this predictor, values were split into groups by the median ρ in the sample.

We used temporal clustering at retrieval and retrieval performance as the two main behavioral outcome variables. Temporal clustering was calculated as previously published *(47)*. In brief, for each retrieval period, all transitions between recalled words were considered. If there were four words, then there were three transitions between them. For example, “queen” to “world” counted as a transition. The difference in encoding serial positions for both words of the transition was then obtained. Each transition received a score based on where the actual transition serial position difference ranked among all possible remaining transitions. Because of the noise of this metric on a single-list level, we computed the metric across the entire session by averaging all transitions made throughout the session. Due to the nature of the metric, only encoding lists from which subjects recalled at least two words contributed.

We used a two-sample Kolmogorov-Smirnov test to assess the similarity between the performance-by-serial position curve and the peak-count-by-serial position curve. We obtained the latter by counting the number of time cell PSTHs that peaked within each of the serial positions. We then normalized both curves to their respective minimum and maximum values to obtain unitless, comparable curves. To control for differences in the performance versus serial position curve between recording sessions in which subjects were presented with 12-word lists and 15-word lists, we considered these sessions separately.

### Statistical Testing

The variables of interest in this study were not generally normally distributed. Therefore, unless otherwise specified, non-parametric tests were used to assess significance. This includes using the rank sum test for comparison of mean rank between two groups, Kruskal-Wallis for comparison of mean rank between three or more groups, and the Spearman rank correlation coefficient for all correlations. For all generalized linear models, we tested predictor coefficients for significance with their associated t-statistics. For unpaired proportions (for example, fraction of time cells versus SME cells whose activity is significantly predicted by encoding success), we used the Chi-square test. We used Fischer’s exact test when less than 80% of the cells in the contingency table of interest had an expected value of less than 5. We used two-tail tests unless otherwise specified. We selected an alpha of 0.05 for all analyses. Where necessary, more details on the statistical testing utilized for a specific analysis can be found under the appropriate heading.

## Supplementary Text

The identification of a significant fraction of time cells with our procedure was not dependent on either the number of time bins used in the Kruskal-Wallis or the integration window of the Gaussian kernel. When using 40 time bins but a Gaussian kernel with a standard deviation of 0.25 s and 0.75 s to convolve the spike train, we isolated 78 and 82 time cells respectively. When using a Gaussian kernel with a standard deviation of 0.5 s but varying the number of bins used in the Kruskal-Wallis to 20 and 80, we isolated 65 and 81 time cells respectively. All yield fractions exceeded that expected by chance (509 candidate cells, P < 0.001, binomial test). Identification of a significant fraction of time cells did not depend on electrodes near epileptogenic tissue. Excluding all electrodes recording inter-ictal activity at some point during implantation, we isolated 33 time cells (33/308 candidate cells, P < 0.001, binomial test). Importantly, we did not include any session with seizure activity or aura.

Our ability to identify time cells significantly depended on the total number of individual units recorded in a given session (r = 0.717, P < 0.001, Spearman rank correlation). This suggests that our ability to isolate time cells largely depended on how many single units we isolated from that session, opposed to from other unidentified factors. As an additional control, we identified a significant fraction of time cells when only using one session from each subject (67/409 candidate cells, P < 0.001, binomial test). For this analysis, we selected the recording session with the highest number of candidate neurons (hippocampal and entorhinal pyramidal cells). In cases of a tie between sessions, we chose the later session.

As an additional control, we identified single units for which firing rate increased following presentation of recalled as compared to non-recalled memory items (“subsequent memory effect” (SME) cells, P < 0.05, rank sum test, Fig. S7). The proportion of time cells that showed a significant SME did not significantly differ from that of the population at large (Yule’s Q = 0.23, P = 0.13, Fig. S8A). We tested firing rate within the preferred time field across time cells and observed no difference between recalled and non-recalled items (Fig. S6B, z(71) = 0.0712, P > 0.2, rank sum test).

We sought to ensure that our identification of time cells was not confounded by sensitivity of firing rate to memory encoding success, appearance of items on the screen, or the semantic content of the items. Adapting control methods used for time cells in rodents *(2)*, we designed a generalized linear model (GLM) with these variables and time field as categorical predictors (Fig. S5A, see supplementary materials). 58 of 79 time cells’ activity significantly depended on time field, significantly more than the number modulated by all other variables (χ^2^(3) = 109.7, P < 0.001). We additionally ran the GLM with all SME cells not concurrently time cells. 50 of 80 SME cells’ activity increases significantly depended on future recall success while only 37 of 80 SME cells’ activity increases significantly depended on time field. Therefore, as expected, SME cells depended on future recall success significantly more than time cells (50/80 versus 16/79, χ^2^(1) = 29.2, P $<$ 0.001) while time cells depended on time field significantly more frequently than SME cells (37/80 versus 58/79, χ^2^(1) = 12.2, P < 0.001).

Regarding time cell precession, only 53% (38/72) of time cells exhibited a negative slope (consistent with precession) irrespective of the significance level of the time-phase relationship, indicating that phase precession occurs in a subgroup of time cells, different than rodent place cells. Still, our findings support the possibility that at least a fraction of time cells may also exhibit place cell-like properties motivating further experimentation in humans (using alternate paradigms) to directly examine this question.

Regarding time cell behavior, we observed that the population of preferred fields of time cells yielded a pattern similar to the serial position curve reliably seen in episodic memory and similar to the serial position curve seen in our participants (Fig. 1D and 1E, 3A) *(48)*. In fact, we found that distribution of peak time cell firing and recall performance across the encoding list did not differ significantly for either sessions with 15-word lists or 12-word lists (P = 0.052 and P = 0.19 respectively, Kolmogorov-Smirnov test, Fig. S12). While time cell firing itself is not significantly modulated by memory encoding success (see Fig. S8B), in aggregate our data show that time cells more accurately represent temporal information at early and late time bins in an encoding list (Fig. S12). This feature of time cells may underlie the human tendency to more strongly remember items presented in these positions.

---

**Fig. S1.**
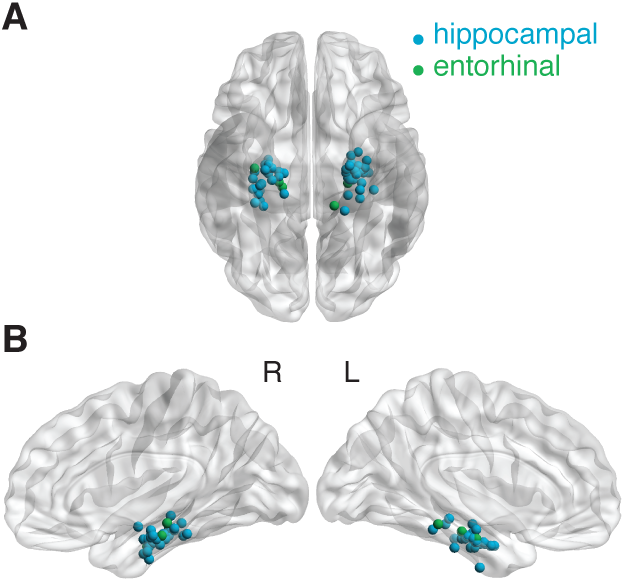
Electrode localizations. (A) Ventral view of the brain with superimposed locations of hippocampal and entorhinal electrodes. (B) Contacts displayed from a medial view.

**Fig. S2.**
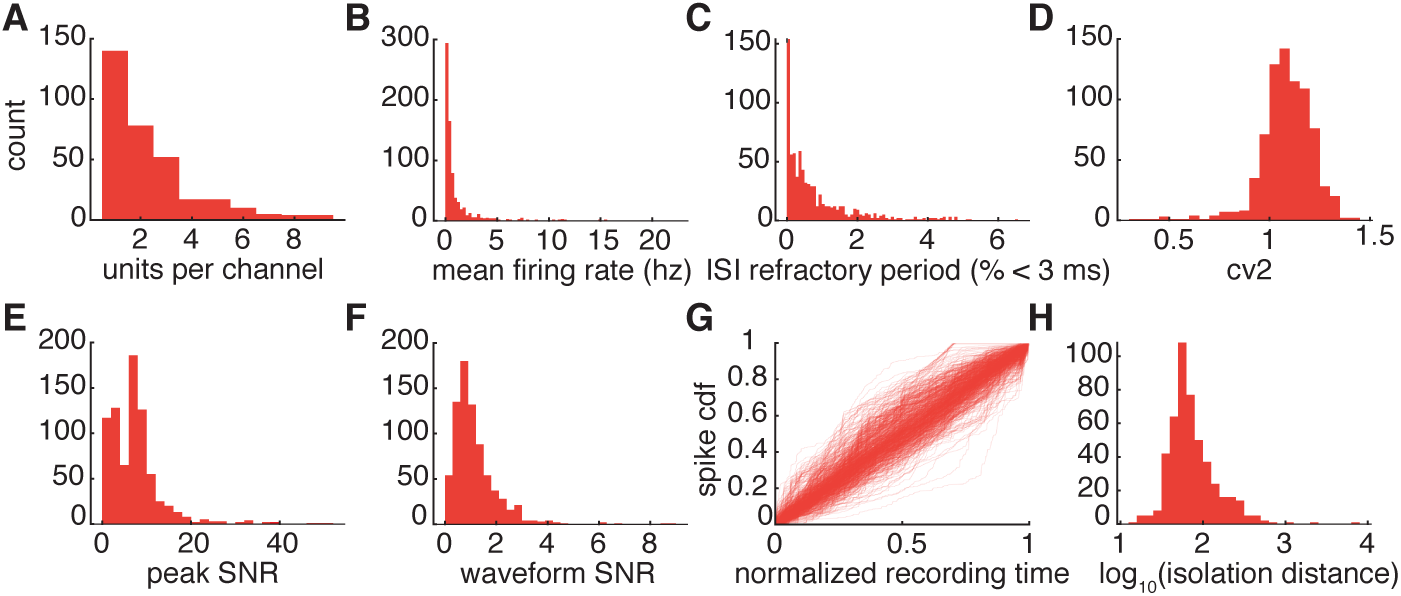
Single unit quality metrics. (A) Histogram of number of units isolated per microelectrode channel. (B) Histogram of mean firing rates of all isolated neurons. (C) Histogram of the percentages of the inter-spike-intervals violating the 3 ms refractory period for each unit. (D) Histogram of CV2 values. (E) SNR histogram, using the peak voltage of the mean spike waveform as the “signal.” (F) SNR histogram, using the mean voltage of the mean spike waveform as the “signal.” (G) Spike cumulative distribution function for each neuron. (H) Histogram of isolation distance values calculated using units isolated from channels recording from at least two units.

**Fig. S3.**
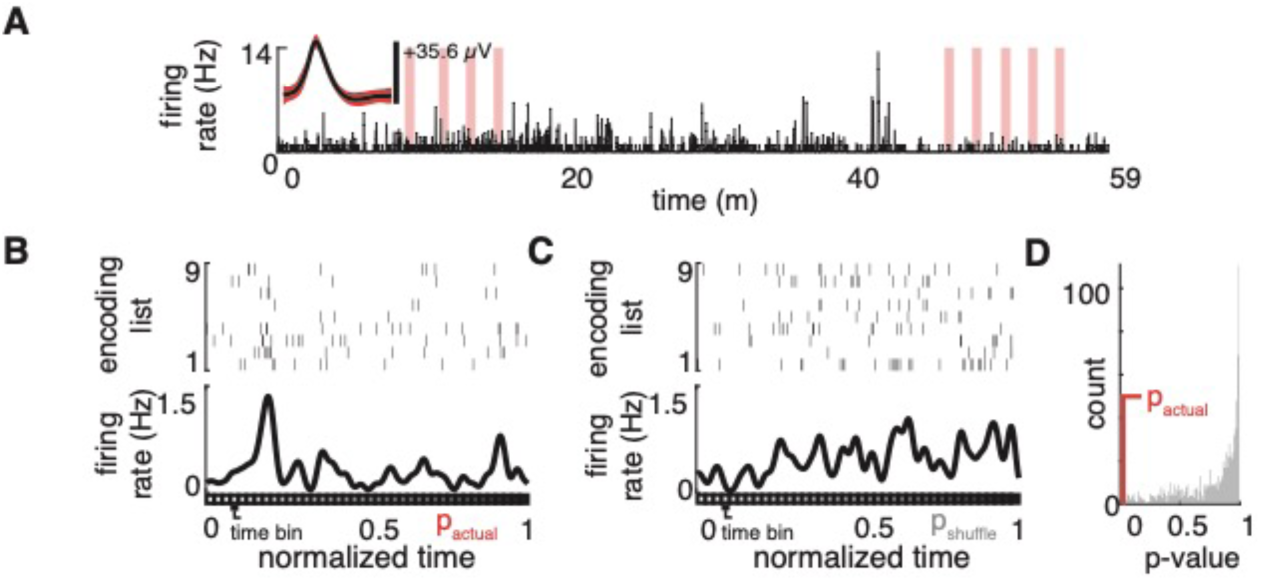
Time cell circular shuffle procedure. (A) An example neuron’s tuning curve across the entire recording session. Its mean waveform is displayed and times representing the session’s encoding lists are highlighted. (B) The actual spike rasters for the example cell from (A) during each encoding list (top) shown with the peri-stimulus time histogram (PSTH) (bottom). Time bins used in the Kruskal-Wallis are displayed along the time axis. (C) The encoding list spike rasters (top) and PSTH (bottom) obtained after circularly shifting the actual tuning curve in (A). (D) Distribution of p-values from 1,000 repetitions of random tuning curve circular shifts.

**Fig. S4.**
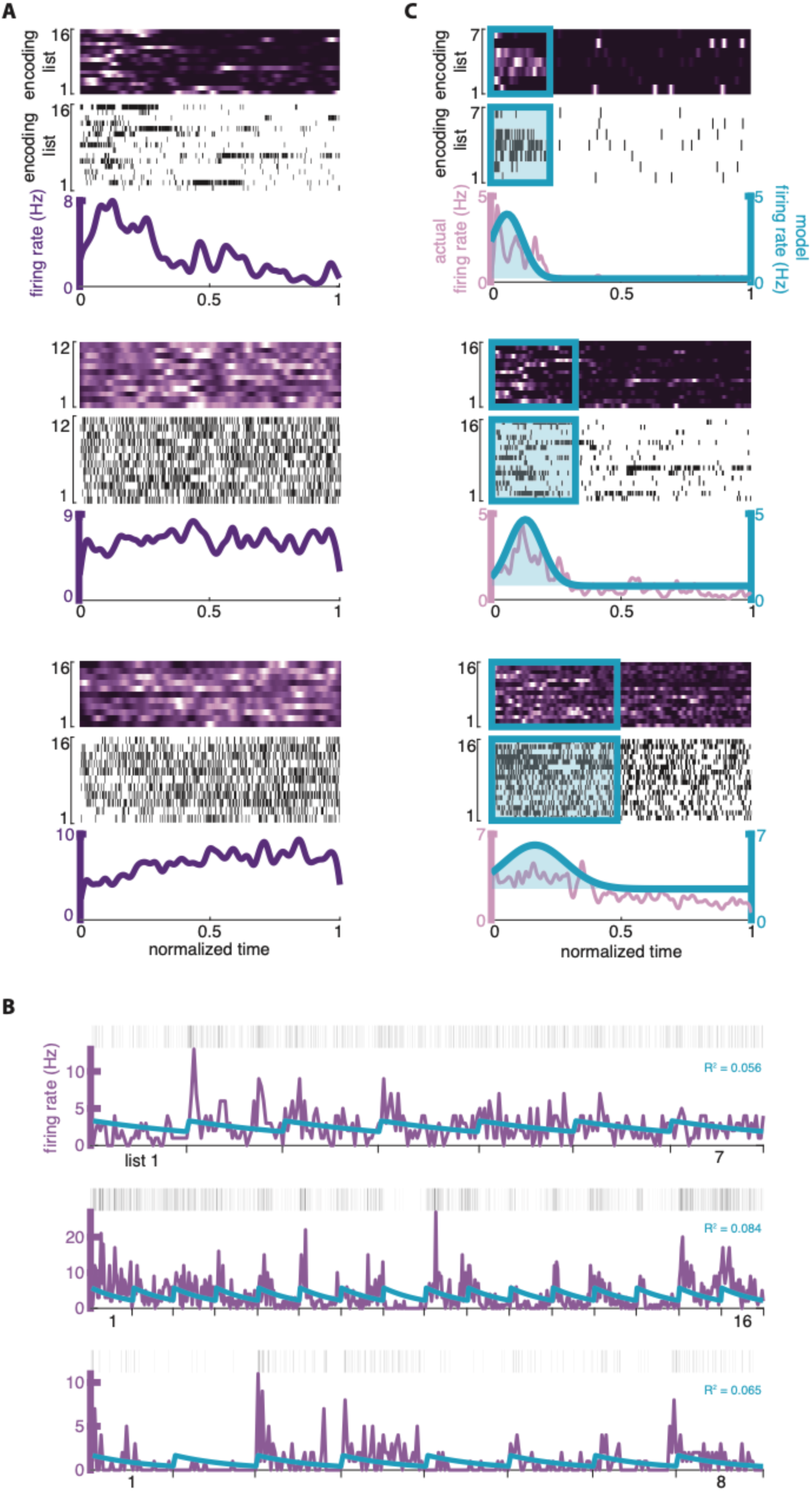
Additional examples of time cells, field cells, and ramping cells. (A) 3 examples of time cells. Spike heat map (top), spike rasters (middle), and PSTH (bottom) are plotted against normalized encoding list time for each cell. (B) 3 examples of time field cells. The spike heat map (top), spike raster plots (middle), and session PSTH with the superimposed model-predicted firing rate (bottom) are displayed for each cell. (C) 3 examples of ramping cells. Juxtaposed encoding list spike rasters (top) and tuning curves with superimposed model-predicted firing rate (bottom) are displayed for each neuron.

**Fig. S5.**
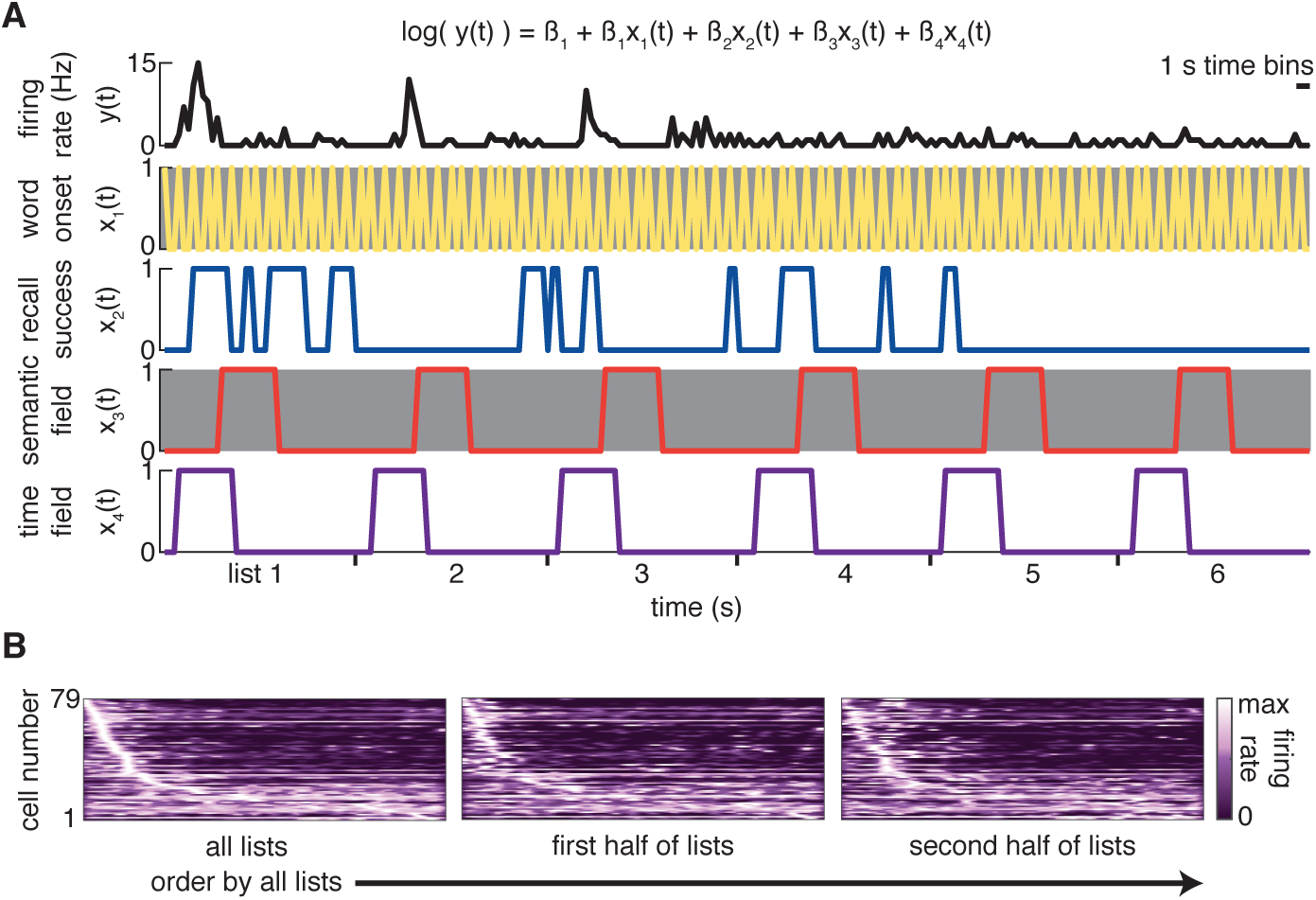
Main control analyses. (A) Design of the control GLM. The outcome variable (firing rate, top) from juxtaposed encoding lists is shown on top of four predictor variables (word onset, recall success, semantic field, time field). (B) PSTH heat maps for all time cells. For all heat maps, rows are ordered by the time sequence of PSTH peaks when considering all encoding lists.

**Fig. S6.**
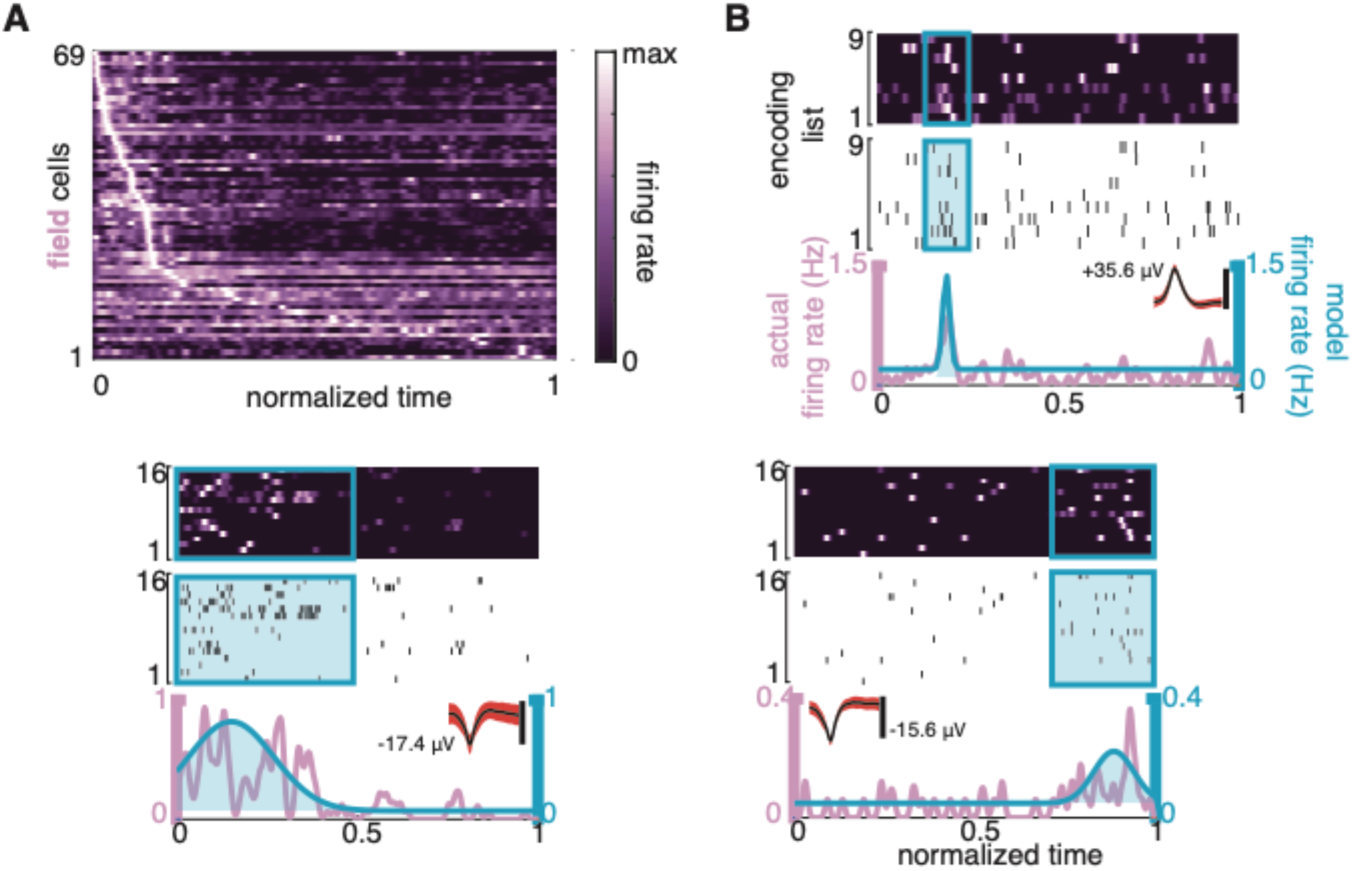
Time field cells. (A) Heat map of all PSTHs from all time field cells, ordered by time of peak activity. (B) 3 examples of time field cells. The spike heat map (top), spike raster plots (middle), session PSTH with superimposed model firing rate prediction (bottom), and mean spike waveform (bottom, inset) are displayed for each cell.

**Fig. S7.**
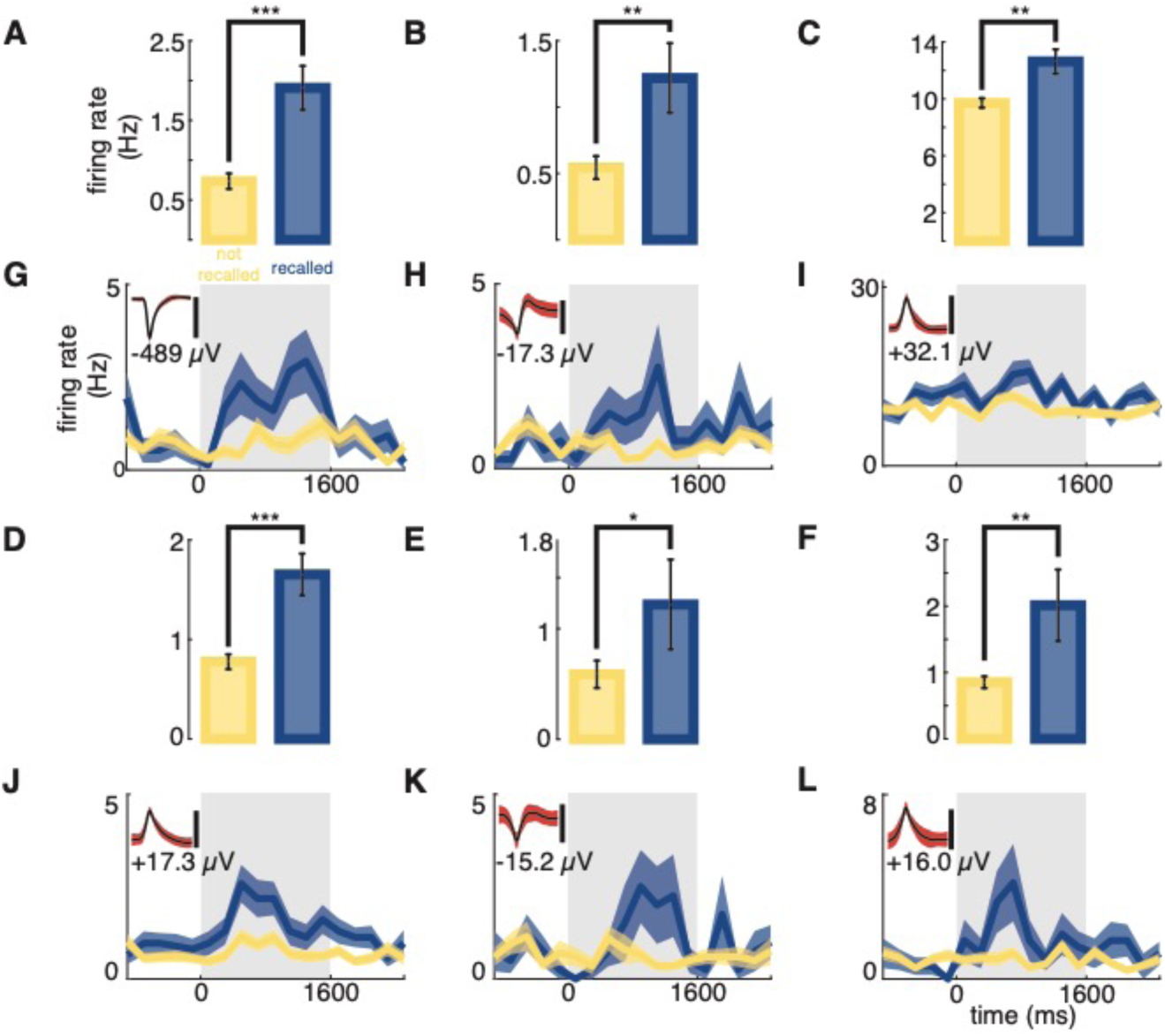
SME cell examples. (A-F) Bar plot showing the average firing rate of the example cell across all recalled (blue) and non-recalled (yellow) encoding periods. Error bars represent the standard error of the mean. *P $<$ 0.05. **P $<$ 0.01. ***P $<$ 0.00001. (G-L) PSTHs for the cells whose data is displayed in (A-F) respectively. Shaded region represents standard error of the mean. The mean spike waveforms with peak voltages of these neurons are inset.

**Fig. S8.**
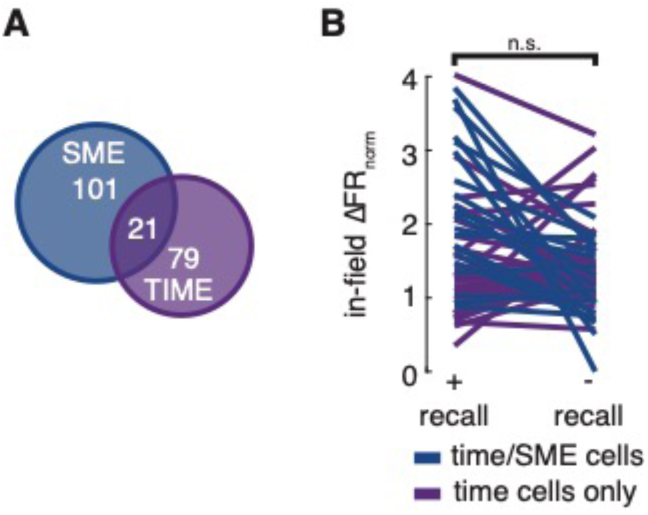
Time cell and SME cell disambiguation (A) Venn-diagram of the overlap between SME and time cell populations. (B) Comparison of in-field firing rate change across all successful versus unsuccessful encoding events across all time cells. n.s. = not significant.

**Fig. S9.**
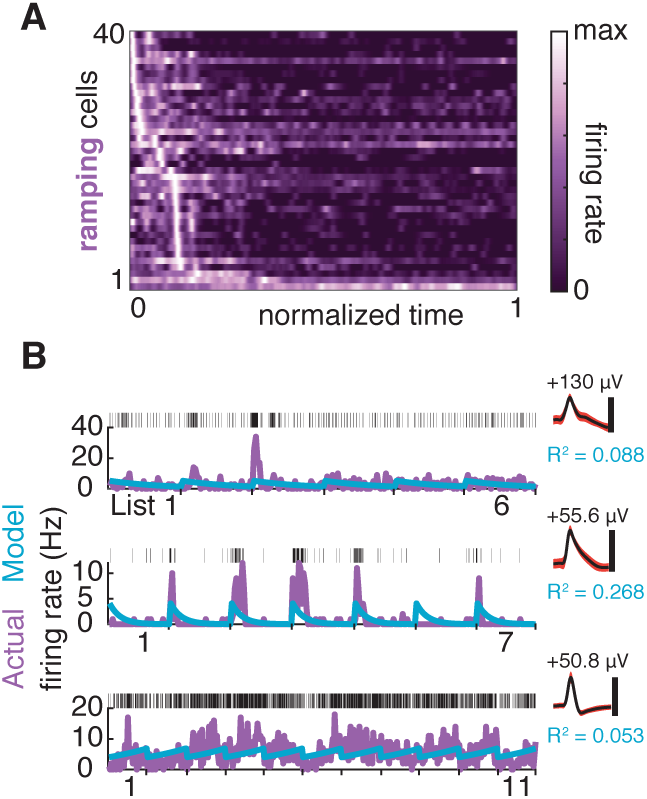
Ramping cells. (A) Heat map of all PSTHs from all ramping cells with an R^2^ of at least 0.05, ordered by time of peak activity. (B) 3 examples of ramping cells. Encoding list tuning curves and spike trains are juxtaposed with intervening data excised. Spike rasters (top), tuning curves with superimposed model-predicted firing rate (bottom), and mean spike waveform (top, right) are displayed for each neuron.

**Fig. S10.**
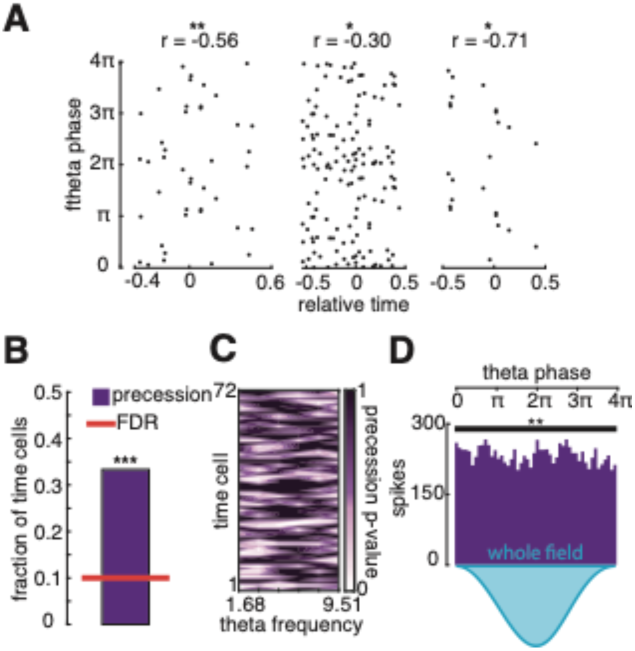
Theta precession and time cells. (A) 3 additional examples of time cell phase-time plots. Data are from the time cells’ peak time field (field with the greatest firing rate). Circular-linear correlation coefficient values are listed above the plots. *P < 0.05. **P < 0.01. (B) Fraction of time cells demonstrating precession after FDR correction with a Q-value of 0.2. ***P $<$ 0.001. (C) Heat map of P-values corresponding to the strength of the association between theta phase and time for each cell at each of 6 log-spaced frequencies. If the slope relating phase and time is positive, the P-value is set to 1 for display purposes only. (D) Phase histogram of all spikes occurring within the preferred time fields (highest peak firing rate) of all time cells with theta precession. **P $<$ 0.01.

**Fig. S11.**
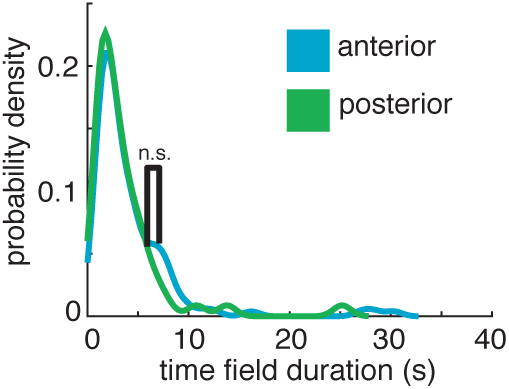
Time field duration along the anterior-posterior hippocampal axis. Comparison of probability density curves for time cell duration between the anterior and posterior hippocampus. n.s. = not significant.

**Fig. S12.**
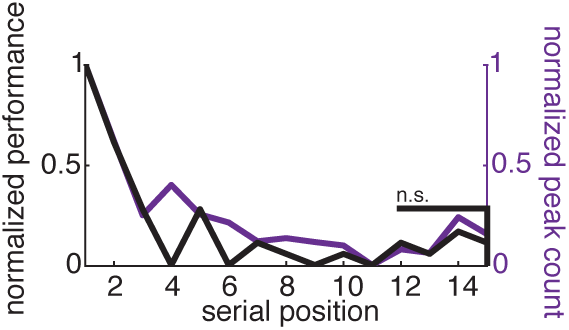
Serial-position curve and time field population curve. Comparison between the performance-by-serial position curve and peak-count-by-serial position curve for all hippocampal time cells extracted from sessions with 15-word lists. n.s. = not significant.

**Table S1.** Abbreviations include: FR = firing rate, SD = standard deviation, ISI = inter-spike-interval, CV2 = modified coefficient of variation, SNR = signal-to-noise ratio, IsoD = isolation distance.

| Neuron count | FR ± SD (Hz) | %ISI < 3 ms ± SD | CV2 ± SD | Peak SNR ± SD | Mean SNR ± SD | Burst Index ± SD | Spike width ± SD | IsoD ± SD |
| --- | --- | --- | --- | --- | --- | --- | --- | --- |
| 768 | 1.34 ± 2.63 | 0.81 ± 1.01 | 1.08 ± 0.13 | 7.39 ± 6.01 | 1.15 ± 0.90 | 0.05 ± 0.05 | 0.52 ± 0.14 | 114 ± 334 |

